# Helical charge distribution at the transmembrane-luminal interface determines subcellular localization

**DOI:** 10.1101/2025.11.12.688035

**Authors:** Haddas Saad, Ron Benyair, Tal Mazor, Gerardo Z. Lederkremer

## Abstract

Transmembrane protein localization is typically dictated by short cytosolic tail sequences, whereas luminal domain contribution remains less understood. Here, we identify a short luminal juxtamembrane peptide as a key determinant of subcellular localization for Class I α-1,2 mannosidases—ERManI, ManIA, ManIB, and ManIC—members of the glycoside hydrolase 47 family involved in N-glycoprotein processing. We previously showed that ERManI and ManIA localize to specialized quality control vesicles (QCVs); we now find that ManIB also partially localizes to these vesicles, whereas ManIC is predominantly Golgi-localized. ERManI, ManIA, and ManIB share a conserved luminal juxtamembrane distribution of charged residues, which diverges in ManIC. Structural predictions suggest that this region maintains an α-helical conformation, with the charge pattern oriented on one face. Site-directed mutagenesis that disrupted this charge pattern or altered its helical register (via alanine insertions) shifted localization between QCVs and the Golgi. Moreover, grafting this peptide onto an unrelated transmembrane protein, β-1,3-galactosyltransferase, redirected it from the Golgi to QCVs. These findings suggest that a specific three-dimensional charge pattern at the transmembrane-luminal interface serves as a localization signal. Unlike canonical linear motifs, this mechanism relies on the structural arrangement, revealing a previously unrecognized mode of organelle targeting within the secretory pathway.

## Introduction

The secretory pathway consists of a series of membrane-bound organelles connected by vesicle-mediated transport and targeted fusion. Membrane and secretory proteins are inserted into the endoplasmic reticulum (ER) where they undergo folding and initial modification before being transported to the Golgi apparatus in COPII-coated vesicles, for complex glycan modifications ^1^. Most Golgi resident enzymes are type II membrane proteins with a small cytoplasmic tail, a single transmembrane domain (TMD), and a luminal catalytic domain ^2, 3^. The TMD plays a critical role in retaining enzymes within the Golgi, and the retention mechanism is conserved across species, including mammals and yeast ^4^. Mutational analysis of TMDs supports the hypothesis that retention is due to physical properties of the TMDs rather than protein-protein interactions ^5^.

Most transmembrane proteins have hydrophobic α-helical TMDs enabling them to partition out of the Sec61 translocon into the ER membrane during synthesis ^6^. A comprehensive analysis using genome data of single TMD proteins demonstrated that TMDs are highly specialized, reflecting distinct physical properties of membranes along the secretory pathway ^7^. The length, charge distribution, hydrophobicity and residual composition of TMDs are key determinants of membrane protein localization. The residue composition and hydrophobic length of a TMD are influenced by the physical properties of the bilayer (e.g., thickness, lipid order), which differ between organelles. Proteins destined for ER residency must carry specific topogenic signals, while those lacking such signals are transported to the plasma membrane by default ^8^. The TMD length was found to play a crucial role in retaining ER-resident proteins; specifically, short TMDs preclude ER tail-anchored proteins from progressing down the secretory pathway, thus, the short membrane anchor serves as a general mechanism for retaining transmembrane ER proteins, preventing their inclusion in vesicular transport beyond the ER, while proteins destined for post-Golgi compartments have longer TMDs ^1, 9^. TMD hydrophobicity has also been found to affect localization, as proteins with low TMD hydrophobicity can integrate directly into lipid bilayers, while more hydrophobic TMDs require chaperones or alternative pathways such as TMD recognition complexes ^10^. Selective signals within the cytoplasmic domains of membrane proteins, such as di-hydrophobic and di-acidic motifs, have been found to be required for efficient transport between organelles. A di-acidic motif (DXE) on the cytosolic C-terminus of Type I membrane proteins was identified as a critical signal for the recruitment of COPII vesicles and the selective export of proteins from the ER to the Golgi apparatus ^11^. Furthermore, in the case of multimeric membrane proteins, arginine-based ER localization signals (RXR) on cytoplasmic domains were found to be essential for their quality control, regulation and trafficking, by retaining improperly assembled protein subunits in the ER until they are correctly assembled into complexes, at which point the signal is masked via steric hindrance, phosphorylation or recruitment of regulatory proteins, allowing export from the ER. These motifs function independently of their position relative to the protein’s termini, distinguishing them from other ER signals such as KXK motifs, which are usually close to the terminus and far from the TMD ^12^. An additional COPII-dependent ER export motif of glycosyltransferases, which are type II membrane proteins residing in the Golgi apparatus, is the di-basic motif (RK(X)RK), on the cytosolic domain adjacent to the TMD. This motif is conserved among different glycosyltransferases in different organisms. Mutations in this motif, such as replacing arginine and lysine with alanine, disrupt Golgi localization and lead to ER retention, and chimeric constructs replacing the cytosolic tails of ER-resident proteins with those of glycosyltransferases can exit the ER ^13^. In contrast to cytosolic signals, luminal motifs are less characterized on transmembrane proteins, although the luminal C-terminal KDEL is the main ER retrieval motif for soluble ER resident proteins ^14^. We had previously determined that a type II membrane protein, an ER-associated degradation (ERAD) substrate, human asialoglycoprotein receptor (ASGPR) precursor H2a ^15^, contains a luminal juxtamembrane pentapeptide, EGHRG, which we identified as signal for ER retention ^16^ and subsequent routing to a specialized juxtanuclear compartment, the ER-derived quality control compartment (ERQC), a staging ground for retrotranslocation and targeting to ERAD ^17–19^.

The Glycoside Hydrolase 47 (GH47) family of enzymes, also known as Class I α-1,2 mannosidases, play important roles in the processing of N-glycans. Three subfamilies of GH47 enzymes have been identified based on their different specifities: ER α-1,2 mannosidase I (ERManI/ MAN1B1), α-1,2 mannosidases IA, IB, and IC (ManIA/ MAN1A1, ManIB/ MAN1A2 and ManIC/ MAN1C1), and the ER degradation-enhancing mannosidase-like (EDEM) proteins ^20^. The α-1,2 mannosidases share a high level of homology and are all type II transmembrane proteins with a typical structure which includes a relatively short cytosolic N-terminus, a transmembrane region, and a long luminal carboxy terminal domain including a stem region and a catalytic domain. We have previously shown that, surprisingly, ERManI resides in novel quality control vesicles (QCVs) at the steady state, and not in the ER as was initially thought, and under ER-stress conditions it accumulates in the ERQC ^21, 22^. Unexpectedly, we found that ManIA also resides in QCVs and not in the Golgi complex as was previously thought ^23^. Thus far, no localization signal has been reported for any of the Class I α-1,2-mannosidases. Here we have determined that the three dimensional charge pattern of the TMD-lumen interface sequence functions as a localization signal of ERManI and mannosidases IA, IB and IC.

## Methods

### Materials

Lipofectamine 2000 transfection reagent was from Invitrogen (Cat#52758). cOmplete Protease Inhibitor Cocktail was from Roche (Cat#11697498001). Bortezomib (Bz) (Cat#179324-69-7), OptiPrep Density Gradient Medium (Cat#D1556), BrefeldinA (Cat#B6542) and other common reagents were from Sigma-Aldrich.

### Antibodies

Mouse anti-HA was from Biolegend (Cat#901514 RRID:AB_2565336), rabbit anti-calnexin (Cat#C4731 RRID:AB_476845) and mouse anti-actin (Cat#A3853 RRID:AB_2765344) were from Sigma-Aldrich, Rabbit anti-GFP was from Santa Cruz Biotechnology (Cat#sc-8334 RRID:AB_641123). Rabbit anti-Cab45 was the one used previously ^24^. Goat anti-mouse IgG-HRP (Cat#111-035-144RRID:AB_2307391) and goat anti-rabbit IgG-HRP (Cat#115-035-166RRID:AB_2338511) were from Jackson-Immuno Research Labs.

### Cell culture, media and transfections

NIH 3T3, U2OS and HEK 293 cells were grown in DMEM supplemented with 10% bovine calf serum at 37°C under 5% CO_2_. Transfections of NIH 3T3 and U2OS cells were carried out using Lipofectamine 2000 transfection reagent. Transfection of HEK293 cells was performed using the calcium phosphate method.

### Plasmids and constructs

ERManI-mCherry: ERMan1 cDNA subcloned into mCherry-N1 expression vector (Clonetech) was described before ^21^. HA-tagged ERManI in pMH was used previously ^22^. ERManI-GFP and ManIA-mCherry were described before ^23^. ManIA-GFP: ManIA cDNA was subcloned into eGFP-N1 expression vector (Clonetech) with a flexible linker (SGGGGS), using Sal1 and HindIII. ManIB-HA and ManIC-HA in the pMH expression vector (Roche Diagnostics, Basel, Switzerland) were kind gifts from A. Herscovics (McGill University, Montreal, Canada). ManIB-mCherry and ManIB-GFP: ManIB cDNA was subcloned into mCherry-N1 or eGFP-N1 expression vectors respectively (Clonetech) with a flexible linker (SGGGGS), using Sal1 and HindIII. ManIC-mCherry and ManIC-GFP were constructed similarly to ManIB constructs but using ManIC cDNA. H2a-GFP: H2a linked to GFP with the flexible linker SGGGGS was described in ^25^. GalT-YFP ^21^ and GalT-CFP (first 60 amino acids of β-1,3-galactosyltransferase linked to YFP or CFP) were kind gifts from K. Hirschberg (Tel Aviv Univ.).

ERManI[D106A]: Exchange of aspartate to alanine at position 106 of ERManI-GFP was carried out by PCR, using the following primers: CTACATCAACTTGGCTGCTCATTGGAAAGCTCTGGCTTTCAGG, CCTGAAAGCCAGAGCTTTCCAATGAGCAGCCAAGTTGATGTAG. Similarly, for ERManI[K109A], ERManI[1Ala], ERManI[2Ala], ERManI[3Ala] using the following primers:

ERManI[K109A]: CTACATCAACTTGGCTGACCATTGGGCTGCTCTGGCTTTCAGG, CCTGAAAGCCAGAGCAGCCCAATGGTCAGCCAAGTTGATGTAG.

ERManI[1Ala]: CTCTTCTACATCAACTTGGCTGCTGACCATTGGAAAGCTCTG, CAGAGCTTTCCAATGGTCAGCAGCCAAGTTGATGTAGAAGAG.

ERManI[2Ala]: CTCTTCTACATCAACTTGGCTGCTGCCGACCATTGGAAAGCTCTG, CAGAGCTTTCCAATGGTCGGCAGCAGCCAAGTTGATGTAGAAGAG.

ERManI[3Ala]: CTCTTCTACATCAACTTGGCTGCTGCCGCTGACCATTGGAAAGCTCTG, CAGAGCTTTCCAATGGTCACGGCAGCAGCCAAGTTGATGTAGAAGAG.

ERManIC: The amino acid sequence YINLADHWKA of ERManI was replaced with the amino acid sequence LLPHSSRLKR of ManIC, using the following primers: CTGCTGCCCCACTCCTCTCGCCTCAAGCGCCTGGCTTTCAGGCTAGAGG, GCGCTTGAGGCGAGAGGAGTGGGGCAGCAGGAAGAGGAGTCCACAGAAA AG MANICER: The amino acid sequence LLPHSSRLKR of ManIC was replaced with the amino acid sequence YINLADHWKA of ERManI, using the following primers: GGGCCCTCTTCTACATCAACTTGGCTGACCATTGGAAAGCTCTCTTCCTG, AGCTTTCCAATGGTCAGCCAAGTTGATGTAGAAGAGGGCCCCGAAGC.

The following constructs were made by three-step PCR reactions using the following primers:

ManIC[H45D]: GCCAAGCTTATGCTCATGAGGAAAGTG and CTTGAGGCGAGAGGAGTCGGGC; GCCCGACTCCTCTCGCCTCAAG and AGAGCGGCCGCGTGTCTGCCCCAGGCTC.

ManIB[10C]: Replacement of the amino acid sequence DSSKHKRFDL of ManIB-HA with HSSRLKRLFL of ManIC: TCTAAGCTTATGACTACCCCAGCCCTG and CAGGAAGAGGCGCTTGAGGCGAGAGGAGTGTGGAAGGAAAAAGAATG; CACTCCTCTCGCCTCAAGCGCCTCTTCCTGGGTTTAGAAGATGTG and AGAGCGGCCGCTCGAACAGCAGGATTAC.

ManIC[10B]: Replacement of HSSRLKRLFL of ManIC-HA with DSSKHKRFDL of ManIB: GCCAAGCTTATGCTCATGAGGAAAGTG and CAAATCAAAGCGTTTGTGTTTTGAAGAGTCGGGCAGCAGGAAGAG; GACTCTTCAAAACACAAACGCTTTGATTTGGCCCCCCGGACCCAG and AGAGCGGCCGCGTGTCTGCCCCAGGCTC.

ManIB[Δ58P]: The deletion of proline from position 58 of ManIB-HA: TCTAAGCTTATGACTACCCCAGCCCTG and TGAAGAGTCAAGGAAAAAGAATGCC; GGCATTCTTTTTCCTTGACTCTTCA and AGAGCGGCCGCTCGAACAGCAGGATTAC.

GalT[R45A]: Exchange of arginine to alanine at position 45 of GalT-YFP was synthesized by TWIST Bioscience (Israel).

GalTER: The replacement of the amino acid sequence RDLSR of GalT-YFP with the amino acid sequence DHWKA of ERManI was synthesized by HY Laboratories LTD (Israel).

### Iodixanol equilibrium sedimentation gradient

Iodixanol gradients were performed similarly to previously described ^24^. Briefly, HEK 293 cells transfected with plasmids as mentioned in the results were washed with PBS and resuspended in a homogenization buffer (0.25M Sucrose, and 10mM HEPES pH7.4). The cells were passed through a 21G needle 5 times before homogenization in a Dounce homogenizer (low clearance pestle, 30 strokes). The homogenates were centrifuged at 1000xg for 10 minutes at 4 °C to remove nuclei and cell debris and the supernatants were loaded on top of an iodixanol gradient (10 to 34%). The gradients were ultra-centrifuged at 24,000 rpm (98,500g, Beckman SW41 rotor) at 4 °C for 16 hours. 1ml gradient fractions were collected from top to bottom and run on SDS-PAGE.

### Immunofluorescence Microscopy

Immunofluorescence was performed as described previously ^23, 26^. Briefly, for live cell microscopy, cells were transfected and grown for 24h on glass bottom 35mm plates. Images were captured at 37°C under 5% CO_2_ using a Leica SP8 laser scanning confocal microscope. For fixation purposes, cells grown for 24 h after transfection on coverslips in 24-well plates were fixed with 3% paraformaldehyde for 30 min, incubated with 50mM glycine in PBS, and left unpermeabilized or permeabilized with 0.5% Triton X-100. After blocking with normal goat IgG in PBS/2% BSA, they were exposed to primary antibody for 60 min, washed and incubated for 30 min with a secondary antibody, followed by washes. Specimens were observed on a Leica SP8 laser scanning confocal microscope or LSM 510 meta microscope (for NIH 3T3 cells in Suppl. figures S3 and S4). ImageJ was used to quantify fluorescence intensity and to calculate Mander’s coefficients (using JACOP) for colocalization studies.

### Bioinformatics analysis

The bioinformatics survey was conducted with the help of M. Schushan and N. Ben Tal (Tel-Aviv University). Multiple sequence alignment comparison was performed using T-COFFEE (Tree based Consistency Objective Function For Alignment Evaluation) software (version 8.97_101117), based on combination of several algorithms including Clustal W. ^27^ TMD predictions were done using the TOPCONS web server (https://topcons.cbr.su.se/pred/) ^28^. Structural predictions were done with the AlphaFold web server https://alphafold.ebi.ac.uk/ ^29^. Edmonson wheel depictions were done with the Pepwheel web server https://www.bioinformatics.nl/cgi-bin/emboss/pepwheel.

### Statistical analysis

The results are expressed as average ±SD or mean ±SEM as indicated. Student’s t-test (two-tailed) was used to compare the averages of two groups. Statistical significance was determined at P<0.05 (*), P<0.01 (**), P<0.001 (***), P<0.0001 (****).

## Results

### Similar charge distribution at the transmembrane-luminal interface of ER mannosidase I and on an ERQC localization signal of an ERAD substrate

ERManI plays a crucial role in ERAD, initiating the trimming of mannose residues from N-linked sugar chains, which determines whether nascent glycoproteins are retained in the ER for further folding attempts or targeted for degradation. At the steady state, ERManI resides in highly dynamic quality control vesicles (QCVs), which show ER-like density, but are distinct from the ER or Golgi apparatus. Only occasionally do glycoproteins interact with ERManI at QCV fusion sites with the ER, resulting in a slow mannose-trimming process ^17, 21^. Under ER-stress conditions, the QCVs converge in the centrosomal region near the nucleus, at the ER-derived quality control compartment (ERQC) ^21^. Examination of H2a fused to GFP in live-cell microscopy showed a very low level of colocalization with ERManI-mCherry (Fig.1a), which could likely be attributed to accelerated degradation of H2a, as we have previously shown that the proteasomal degradation of H2a is accelerated in presence of ERManI ^22^, while following proteasomal inhibition with bortezomib (Bz), both proteins accumulate in a region that we previously characterized as the ERQC ^21^ and show a high level of colocalization (Fig.1b). As we had seen before, this region does not colocalize with a Golgi marker (Fig. 1a, b).

**Fig.1.**
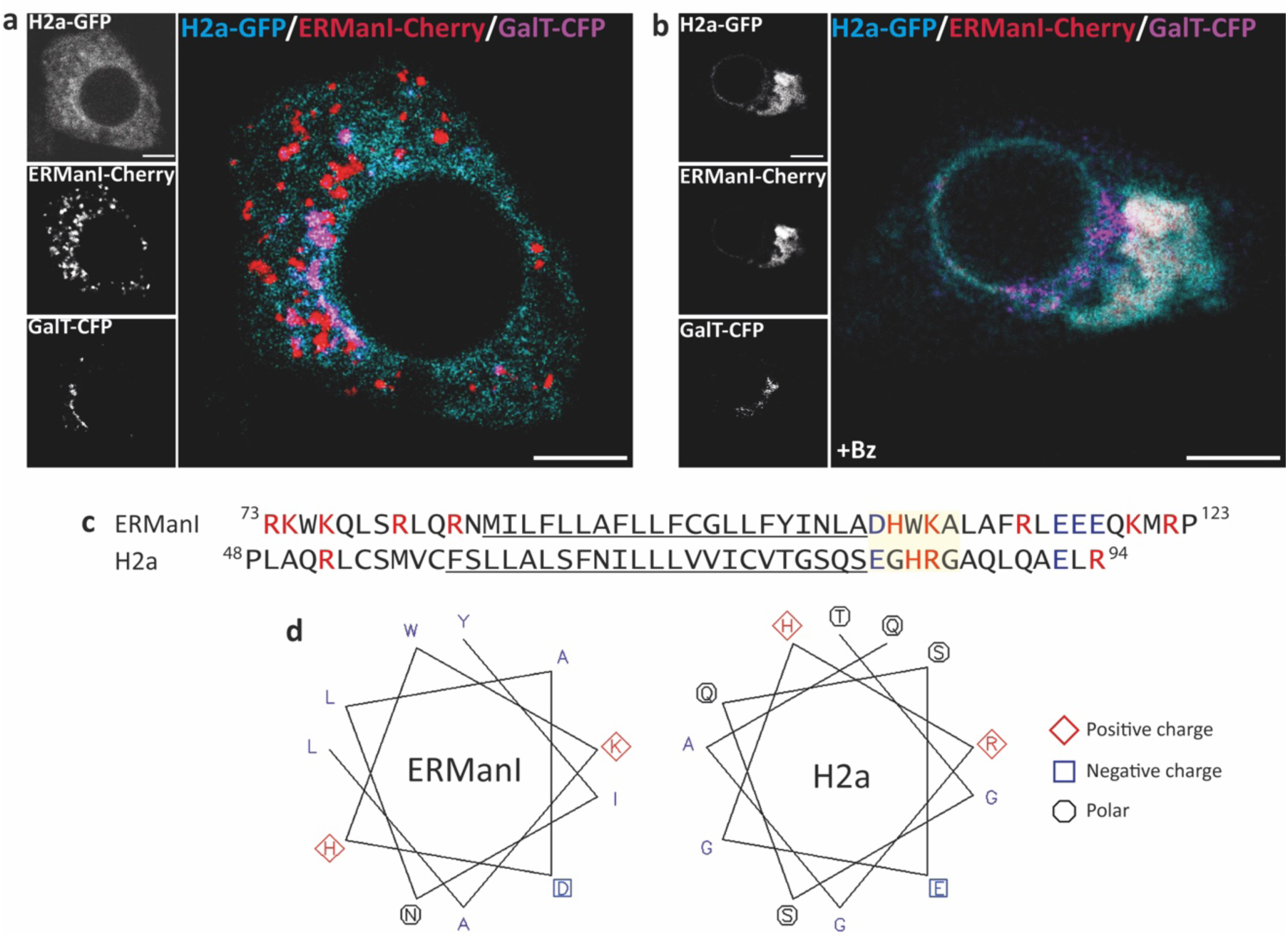
A similar juxtamembrane luminal charge distribution between ERManI and H2a. (a, **b)** NIH 3T3 cells were transfected with plasmids expressing ERManI-mCherry together with H2a-GFP and GalT-CFP (Golgi marker) and observed live 21 hours post transfection. The cells were **(a)** left untreated or **(b)** treated with Bz (1µM) for 3h, and images were captured on a confocal microscope. H2a-GFP, ERManI-Cherry and GalT-CFP have been pseudo-colored cyan, red and magenta, respectively, for better visualization. Colocalization appears as white in the merged images in this and all other figures. Bars=10μM. **(c)** Sequence alignment of human ERManI and H2a. Positively charged amino acids are in red and negatively charged in blue. The juxtamembrane luminal pentapeptide with conserved charge is highlighted in yellow. **(d)** Edmonson alpha helix wheels of the luminal-transmembrane interfaces of ERManI and H2a.

We compared the luminal juxtamembrane region of ERManI with that of H2a, where the charged pentapeptide signal for ERQC targeting resides. This sequence of ERManI has a similar distribution of charged amino acids as in H2a, specifically a negative charge at position 1 after the TMD and a positive charge at position 4 (Fig. 1c), which is conserved among most ERManI orthologues (Suppl. Fig. S1). This region is predicted to preserve the alpha helix structure of the TMD. When comparing the Edmonson alpha helix wheels of ERManI and H2a in the TMD-luminal interface, both helices display a similar distribution of these two charged residues (Fig. 1d). Other regions of the helix are more polar for H2a, which would be expected for an ERAD substrate^30, 31^.

### Comparison with mannosidases IA, IB and IC

We have previously reported that ManIA is not located in the Golgi but resides in QCVs, similar to those carrying ERManI ^23^. Given that ManIA-GFP shows high colocalization with ManIA-cherry, the fluorescent protein that is fused does not appear to have a detectable influence (Fig. 2a, g). When examining GFP fused ERManI in live cell microscopy, there is a high level of colocalization (although not complete) with ManIA-mCherry in a punctate pattern (QCVs) (Fig. 2b, g). While a part of the population of ManIB-GFP also appears in a punctate pattern, it shares a slightly lower level of colocalization with ManIA-mCherry (Fig. 2c, g). The colocalization is much lower between ManIA-cherry and ManIC-GFP, indicating that they reside in different types of vesicles (Fig. 2d, g). In addition, a portion of ManIB and especially ManIC molecules that do not colocalize with ManIA appear in a juxtanuclear pattern. These juxtanuclear molecules colocalize with a Golgi marker (Fig. 2e, f, h).

**Fig.2.**
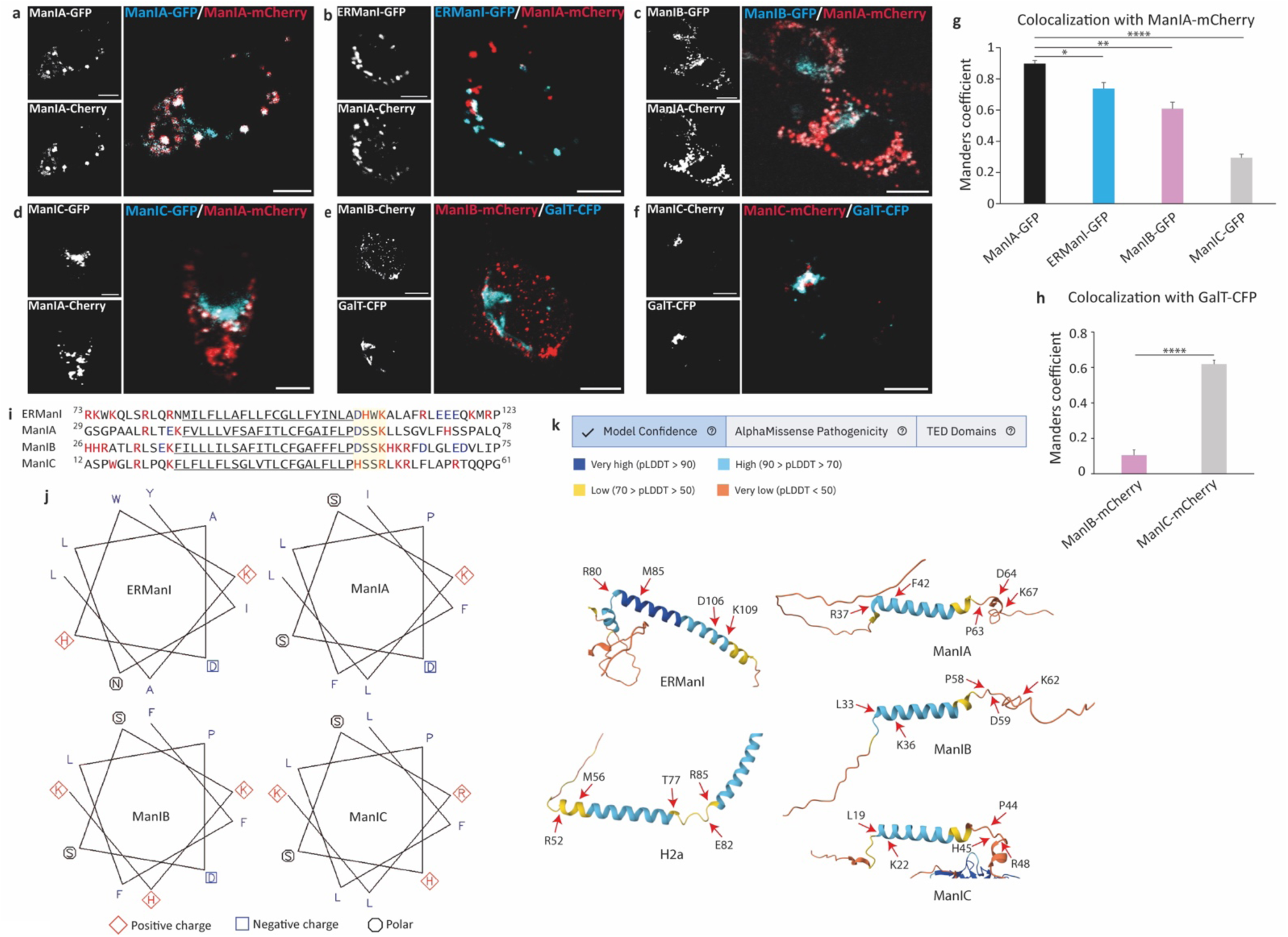
Juxtamembrane luminal charge distribution of ManIA, ManIB, ManIC and ERManI correlates with their QCV or Golgi localization. **(a-f)** NIH 3T3 cells were transfected with plasmids expressing ManIA-mCherry together with (a) ManIA-GFP or (b) ERManI-GFP or (c) ManIB-GFP or (d) ManIC-GFP or GalT-CFP together with (e) ManIB-mCherry or (f) ManIC-mCherry and observed live at 24 hours post transfection on a confocal microscope. GalT-CFP has been pseudo-colored cyan. Bars=10μM. **(g, h)** Graphs of Mander’s coefficients indicating colocalization between each construct with ManIA-cherry (n=17, *P* value (ERManI-GFP)=0.043, P value (ManIB-GFP) = 4.37 · 10^−3^, *P* value (ManIC-GFP) = 2.31 · 10^−11^), or of ManIB and ManIC with GalT-CFP (n=17, *P* value = 3.26 · 10^−11^). **(i)** Sequence alignment of the predicted transmembrane domain (underlined), and flanking regions of human ERManI, ManIA, ManIB and ManIC. **(j)** Edmonson alpha helix wheels of the luminal-transmembrane interface of the mannosidases. **(k)** Predicted structure of the transmembrane domain and flanking regions of the mannosidases and H2a by AlphaFold, highlighting selected residues. Color coding is by model confidence in each region, as indicated.

Sequence comparison of ERManI and mannosidases IA, IB and IC, taking into account many orthologs, shows two regions of high similarity, the extended luminal catalytic region and the TMD and its flanking sequences (Suppl. Fig. S2). There is much less homology in the cytosolic tail and the luminal stem regions. Given the importance of the TMD and flanking sequences for transmembrane protein localization and topology^32, 33^, and the high degree of conservation between these mannosidases, we focused on this region to try to identify a localization signal. The charge distribution at the juxtamembrane cytosolic region is similar between ERManI and ManIC, and between ManIA and ManIB (Suppl. Fig. S1). In contrast, the charge distribution at the juxtamembrane luminal region is similar between ERManI, ManIA and ManIB, but different in ManIC. This pattern at the TMD-luminal interface correlates well with the subcellular localization of the mannosidases, which differs for ManIC (Fig. 2a-h). Therefore, we focused on the TMD-luminal interface of the mannosidases. There are some similarities but also distinct differences in charge distribution (Fig. 2i). Whereas ERManI, ManIA and ManIB share a similar charge distribution, with a negative charge at position 1 after the TMD and a positive charge at position 4, the negative charge is absent in the sequence of ManIC, where it is replaced by a positive charge (Fig. 2i). In addition, another positive charge present in position 2 in ERManI is absent in the other mannosidases. Edmonson helix wheels highlight the similar orientation of the lysine in position 4 and the aspartate in position 1, replaced by histidine in ManIC (Fig. 2j). These residues are oriented on one face of the helix. Therefore, a similarly oriented negative charge in position 1 and positive charge in position 4 correlate with QCV localization (ERManI, ManIA, ManIB), whereas replacement of the negative charge with a positive one correlates with increased Golgi localization (ManIC). AlphaFold predicts with relatively high confidence a continuation of the TMD alpha helix at the juxtamembrane luminal region for ERManI, but with lower confidence and lower helical character in H2a and in ManIA, ManIB and ManIC (Fig. 2k).

### Alteration of charge distribution relocates ERManI to the Golgi

Juxtamembrane charge distribution has a significant effect on protein localization through membrane binding, subcellular compartmentalization, trafficking and protein-protein interactions ^34, 35^. These processes are often affected by electrostatic forces, which are sensitive to changes in the charge distribution of the protein or its local environment. To test the importance of individual charges in the TMD-lumen interface for the localization of ERManI, we exchanged the negatively charged aspartate or the positive charged lysine of the pentapeptide for an uncharged alanine residue, generating the mutants ERManI[D106A] and ERManI[K109A] respectively (Fig. 3a, Table 1). We performed fluorescence microscopy experiments in live NIH 3T3 cells in order to compare the subcellular localization of these mutants fused to GFP with that of WT ERManI fused to mCherry, using β-1,3 galactosyltransferase (GalT) fused to CFP as a Golgi marker. We observed that whereas ERManI-mCherry shows high colocalization with ERManI-GFP, it does not colocalize with ERManI[D106A]-GFP, and hardly colocalizes with ERManI[K109A]-GFP, while both mutants share a high level of colocalization with GalT-CFP (Fig. 3b-f). Density based separation of the different subcellular compartments using iodixanol equilibrium sedimentation gradients confirmed that ERManI[D106A]-GFP and ERManI[K109A]-GFP indeed migrate mostly in the Golgi fractions, while WT ERManI migrates at heavier densities, in ER/QCV fractions as we had seen before ^21^ (Fig. 3g). These results indicate that both the negative charge at position 1 after the TMD and the positive charge at position 4 in the TMD-lumen interface of ERManI have significant effects on its localization.

**Fig.3.**
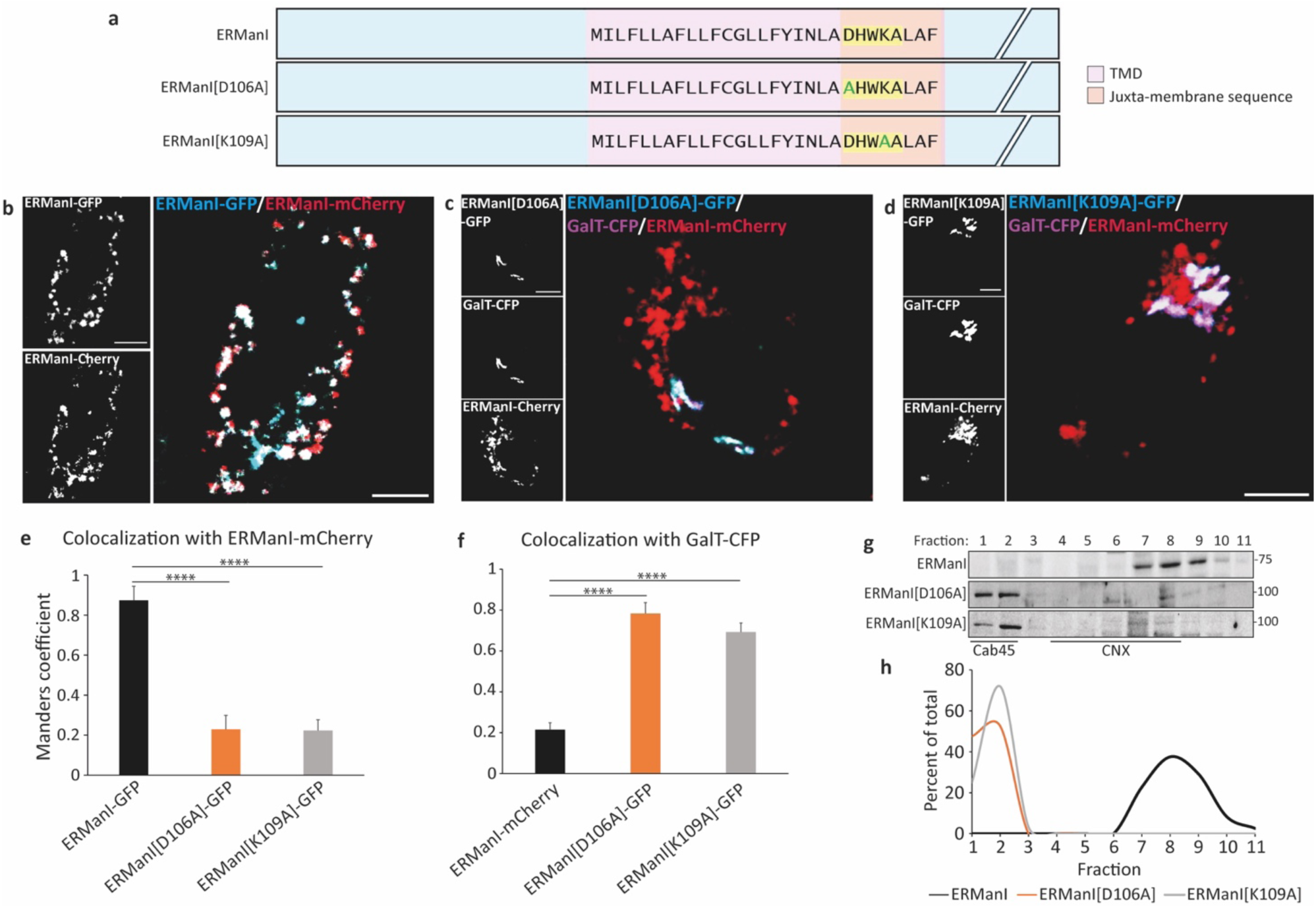
Juxtamembrane luminal charges determine ERManI localization to QCVs or Golgi. **(a)** Schemes illustrating the sequences in the transmembrane and juxtamembrane regions of ERManI and the mutants ERManI[D106A] and ERManI[K109A], with the mutations highlighted in green. **(b-d)** NIH 3T3 cells were transfected with plasmids expressing ERManI-mCherry together with **(b)** ERManI-GFP or **(c)** GalT-CFP and ERManI[D106A]-GFP or **(d)** GalT-CFP and ERManI[K109A]-GFP and observed live at 24 hours post transfection. GalT-CFP has been pseudo-colored magenta. Bars=10μM. **(e, f)** Graphic representation of colocalization of each transfected construct with **(e)** ERManI-cherry (n=13, *P* value (ERManI[D106A]-GFP) = 9.24 · 10^−9^, *P* value (ERManI[K109A]-GFP) = 2.86 · 10^−8^), or **(f)** GalT-CFP (n=13, *P* value (ERManI[D106A]-GFP) = 6.31 · 10^−7^, *P* value (ERManI[K109A]-GFP) = 4.11 · 10^−8^), as inferred by Mander’s coefficient. **(g)** HEK 293 cells were transfected with plasmids expressing ERManI-HA or ERManI[D106A]-GFP or ERManI[K109A]-GFP. 24h post transfection, the cells were homogenized, the homogenate was loaded on top of an iodixanol gradient (10%-34%) and ultracentrifuged as indicated in Methods. Eleven fractions were collected from top to bottom, run on 10% SDS-PAGE and immunoblotted with anti-HA and anti-GFP antibodies. **(h)** The intensity of each band was quantified by ImageJ and plotted as a percent of the total intensity of the protein along the gradient.

**Table 1.** List of constructs. A list of all constructs created and used in the characterization of their localization signal, specification of mutations and purpose.

| **Mutant** | **Mutation** | **Purpose** |
| --- | --- | --- |
| ERManI[D106A] | Exchange of the negatively charged Aspartate of the pentapeptide for an uncharged Alanine residue | Neutralization of one of the two charged amino acids of the pentapeptide to ascertain its role in the localization signal |
| ERManI[K109A] | Exchange of the positively charged Lysine of the pentapeptide for an uncharged Alanine residue | Neutralization of one of the two charged amino acids of the pentapeptide to ascertain its role in the localization signal |
| ERManI[1Ala] | Insertion of an Alanine at location 106 of ERManI | Causes a ~100° shift in helical pitch, changing the orientation of the charged pentapeptide in relation to the transmembrane helix |
| ERManI[2Ala] | Insertion of 2 Alanines at locations 106 and 107 of ERManI | Causes a ~200° shift in helical pitch, changing the orientation of the charged pentapeptide in relation to the transmembrane helix |
| ERManI[3Ala] | Insertion of 3 Alanines at locations 106-108 of ERManI | Causes a ~300° shift in helical pitch, changing the orientation of the charged pentapeptide in relation to the transmembrane helix |
| ERManIC | Replacement of 10 Amino acids in the transmembrane – lumen interface of ERManI with those of ManIC | Implicates the transmembrane – luminal interface region in determination of localization |
| MANICER | Replacement of 10 Amino acids in the transmembrane – lumen interface of ManIC with those of ERManI | Implicates the transmembrane – luminal interface region in determination of localization |
| ManIC[H45D] | Replacement of the positively charged Histidine with the negatively charged Aspartic Acid in the juxta-transmembrane domain of ManIC | Change of the charged amino acid to achieve a pattern similar to that of ERManI |
| ManIC[10B] | Replacement of 10 Amino acids in transmembrane – lumen interface of ManIC with those of ManIB | Implicates the transmembrane – luminal interface region in determination of localization |
| ManIB[10C] | Replacement of 10 Amino acids in transmembrane – lumen interface of ManIB with those of ManIC | Implicates the transmembrane – luminal interface region in determination of localization |
| ManIB[Δ58P] | Deletion of a proline amino acid at position 58 of ManIB | Preserves continuity of the trans and luminal helices |
| GalT[R45A] | Exchange of the positively charged Arginine for an uncharged Alanine residue | Neutralization of the charged amino acids to ascertain its role in the localization signal |
| GalTER | Replacement of 5 Amino acids in GalT with those of ERManI | Implicates the transmembrane – luminal interface region in determination of localization |

We have reported that cell permeabilization with triton-X-100 leads to artificial ERManI localization to the Golgi due to membrane disturbance, emphasizing the need for live-cell imaging ^21^. However, fixation without permeabilization leads to similar results as live cell imaging ^26^. Indeed, also under these conditions ERManI-mCherry colocalizes with ERManI-GFP and does not colocalize with ERManI[D106A]-GFP or ERManI[K109A]-GFP, which appear in a Golgi pattern (Suppl. Fig. S3).

### Changing the orientation of charges by alanine insertion relocates ERManI to the Golgi

Next, we wondered whether the orientation of the charges in respect to the transmembrane helix, as can be observed in the Edmonson helix wheel, is important for ERManI localization. To test this, we used alanine insertion analysis ^36, 37^. We inserted either one alanine residue at position 106, which is predicted to cause a ∼100°shift in the helical pitch, two alanine residues at positions 106 and 107, causing a ∼200°shift, or three alanine residues at positions 106-108, causing a shift of ∼300°, generating the mutants ERManI[1Ala], ERManI[2Ala] and ERManI[3Ala], respectively (Fig. 4a, Table 1). ERManI-mCherry did not colocalize with ERManI[1Ala]-GFP, only slightly colocalized with ERManI[2Ala]-GFP, while it colocalized to a significant extent with ERManI[3Ala]-GFP, as shown by live cell imaging (Fig. 4b-e). ERManI[1Ala]-GFP, and ERManI[2Ala]-GFP showed a high level of colocalization with GalT-CFP (Fig. 4b, c, f). Similar results were obtained in fixed, unpermeabilized NIH 3T3 cells and also in another cells type, in live U2OS cells (Suppl. Fig. S4). Furthermore, iodixanol equilibrium sedimentation gradients confirmed that the insertion of one or two alanine residues led to a complete shift to the lighter fractions, while in the case of three alanine insertions, most of the protein migrated at the same fractions as WT ERManI (Fig. 4g, h). Thus, the orientation of charges appears to substantially influence ERManI localization, shifting it to the Golgi. Insertion of 3 alanines causes a shift in helical pitch of close to 360°, returning the charges to the original orientation on one side of the helix and restoring ERManI localization to a QCV pattern.

**Fig.4.**
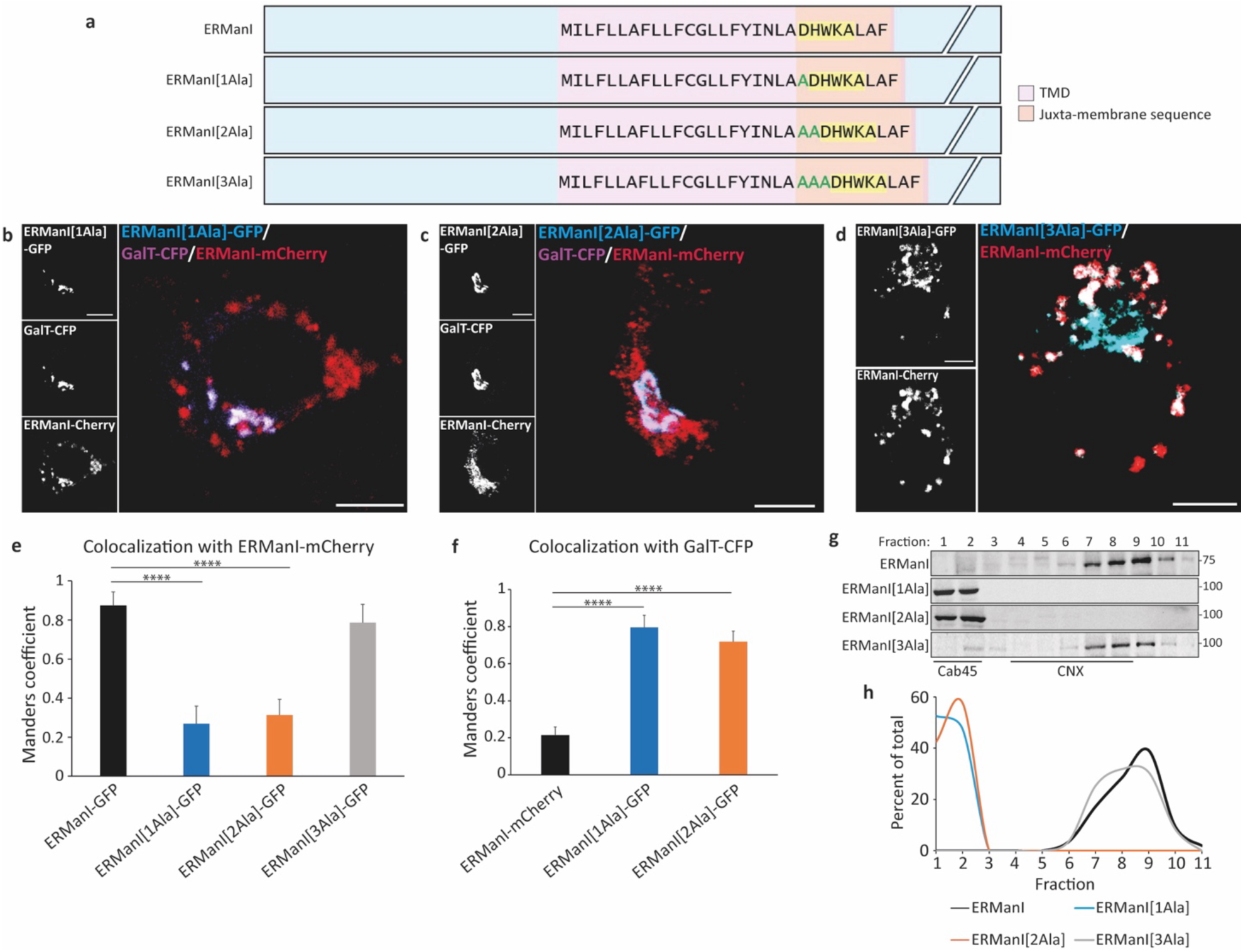
Helical orientation of juxtamembrane luminal charges determine ERManI localization to QCVs or Golgi. **(a)** Schemes illustrating the sequences in the transmembrane and juxtamembrane regions of ERManI and the mutants ERManI[1Ala], ERManI[2Ala] and ERManI[3Ala], with 1 to 3 Ala insertions. **(b-d)** NIH 3T3 cells were transfected with plasmids expressing ERManI-mCherry together with GalT-CFP and **(b)** ERManI[1Ala]-GFP or **(c)** ERManI[2Ala]-GFP or **(d)** ERManI[3Ala]-GFP and observed live at 24 hours post transfection. GalT-CFP has been pseudo-colored magenta. Bars=10μM. **(e, f)** Graphic representation of colocalization of each transfected construct with **(e)** ERManI-cherry (n=12, *P* value (ERMan1ala-GFP) = 2.63 · 10^−7^, *P* value (ERMan2ala-GFP) = 3.35 · 10^−6^), or **(f)** GalT-CFP (n=12, *P* value (ERMan1ala-GFP) = 1.64 · 10^−6^, *P* value (ERMan2ala-GFP) = 6.21 · 10^−6^), as inferred by Mander’s coefficient. **(g)** HEK 293 cells were transfected with plasmids expressing ERManI-HA or the Ala insertion mutants linked to GFP. The cells were homogenized, and the homogenates run on iodixanol gradients as in Fig. 3. **(h)** The intensity of each band was quantified by ImageJ and plotted as a percent of the total.

### The juxtamembrane localization signal can be transferred between ERManI, ManIB and ManIC

Given the distinctly different localization of ERManI and ManIC, we tested whether replacement of the juxtamembrane luminal sequence of ManIC for that of ERManI and vice versa could shift the localization (Fig. 5a, Table 1). Notably, in order to ensure that it is not the TMD length but rather the sequence which influences localization, we intentionally slightly elongated the TMD of ManICER and shortened the TMD of ERManIC, as it is known that longer TMDs are associated with Golgi localization, and shorter TMDs serve as an ER retention signal ^1, 9, 38^. Replacing a ten amino acid segment in the juxtamembrane luminal region of ManIC with the corresponding segment of ERManI (ManICER) resulted in a significant part of the population of ManICER appearing in a vesicular pattern (Fig. 5b). The correlative replacement of this segment in ERManI (ERManIC) led to an accumulation of ERManIC in the Golgi, colocalizing to a high extent with wildtype ManIC-GFP (Fig. 5c, d). The changed shift of these mutants in the density gradients was consistent with the Golgi localization of ERManIC and ER/QCV localization of ManICER, partially comigrating with ERManI (Fig 5e, f).

**Fig.5.**
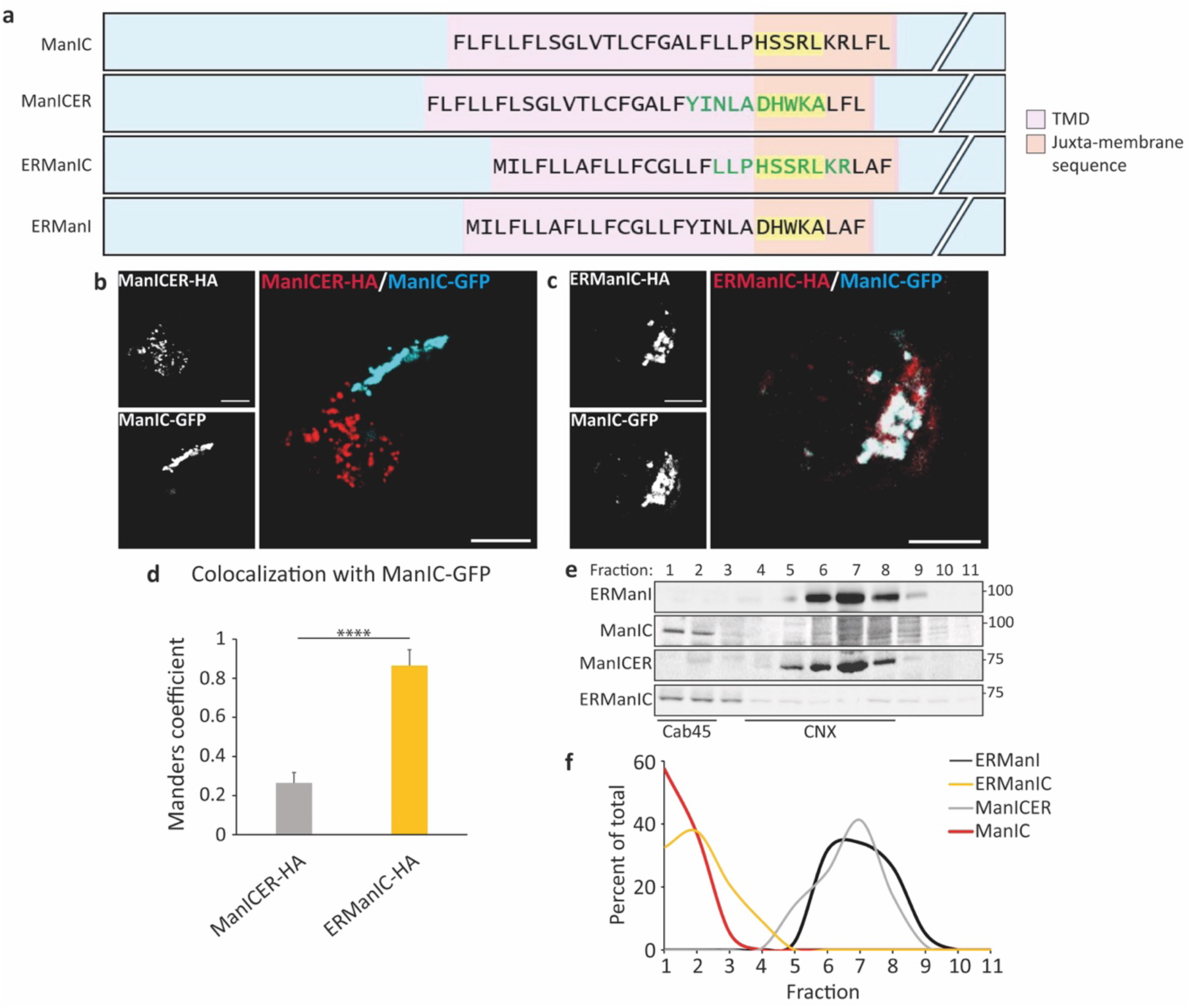
The juxtamembrane luminal peptide determines the QCV or Golgi localization of ERManI and ManIC. **(a)** Schemes illustrating the sequences in the transmembrane and juxtamembrane regions of ERManI and ManIC, and the mutants ERManIC and ManICER, with the luminal juxtamembrane segments swapped. **(b, c)** NIH 3T3 cells were transfected with ManIC-GFP together with (b) ManICER-HA or (c) ERManIC-HA. 24 hours post transfection, the cells were fixed with 3% PFA, permeabilized with 0.5% Triton, subjected to immunofluorescent staining with mouse anti-HA and Dylight594-conjugated goat anti-mouse IgG and imaged on a confocal microscope. Bars=10μM. **(d)** The graph represents the colocalization of each mutant with ManIC-GFP as inferred by Mander’s coefficient. (n=17, *P* value = 6.04 · 10^−12^) **(e)** HEK 293 cells were transfected with plasmids expressing ERManI-GFP or ManIC-GFP or ManICER-HA or ERManIC-HA. The cells were homogenized, and the homogenates run on iodixanol gradients as in Fig. 3. **(f)** The intensity of each band was quantified by ImageJ and plotted as a percent of the total.

We then tested whether exchange of this same segment between ManIB and ManIC can change their localization. Fluorescence microscopy shows that most of the population of ManIB-mCherry appears in a punctate pattern, colocalizing to a relatively low extent with ManIC-GFP, which is mostly concentrated in the Golgi with only a minor part of its population in vesicles (Fig. 6b). Replacement of a ten amino acid segment in the transmembrane-lumen interface of ManIC with the corresponding segment of ManIB resulted in relocation of the mutant ManIC[10B]-HA to vesicles, and the reciprocal exchange led to an increase in the level of colocalization between the mutant ManIB[10C]-HA and ManIC-GFP in the Golgi region (Fig. 6a, c-f, table 1). The shift of the mutants in the density gradients was in line with this change, the migration of ManIB[10C] shifted to the light fractions, whereas ManIC[10B]-HA shifted to heavier fractions, partially comigrating with ManIB (Fig. 6g). To test the importance of a negative charge in position 1 after the TMD (Fig. 2), we replaced the histidine present in ManIC with an aspartic acid, as in the other mannosidases. This replacement in ManIC was sufficient to completely change the localization of the mutant ManIC[H45D], to a punctate pattern with very low colocalization with ManIC-GFP (Fig. 6h-j). Consistently, there was a complete shift in a density gradient to heavier ER/QCV fractions (Fig. 6k).

**Fig.6.**
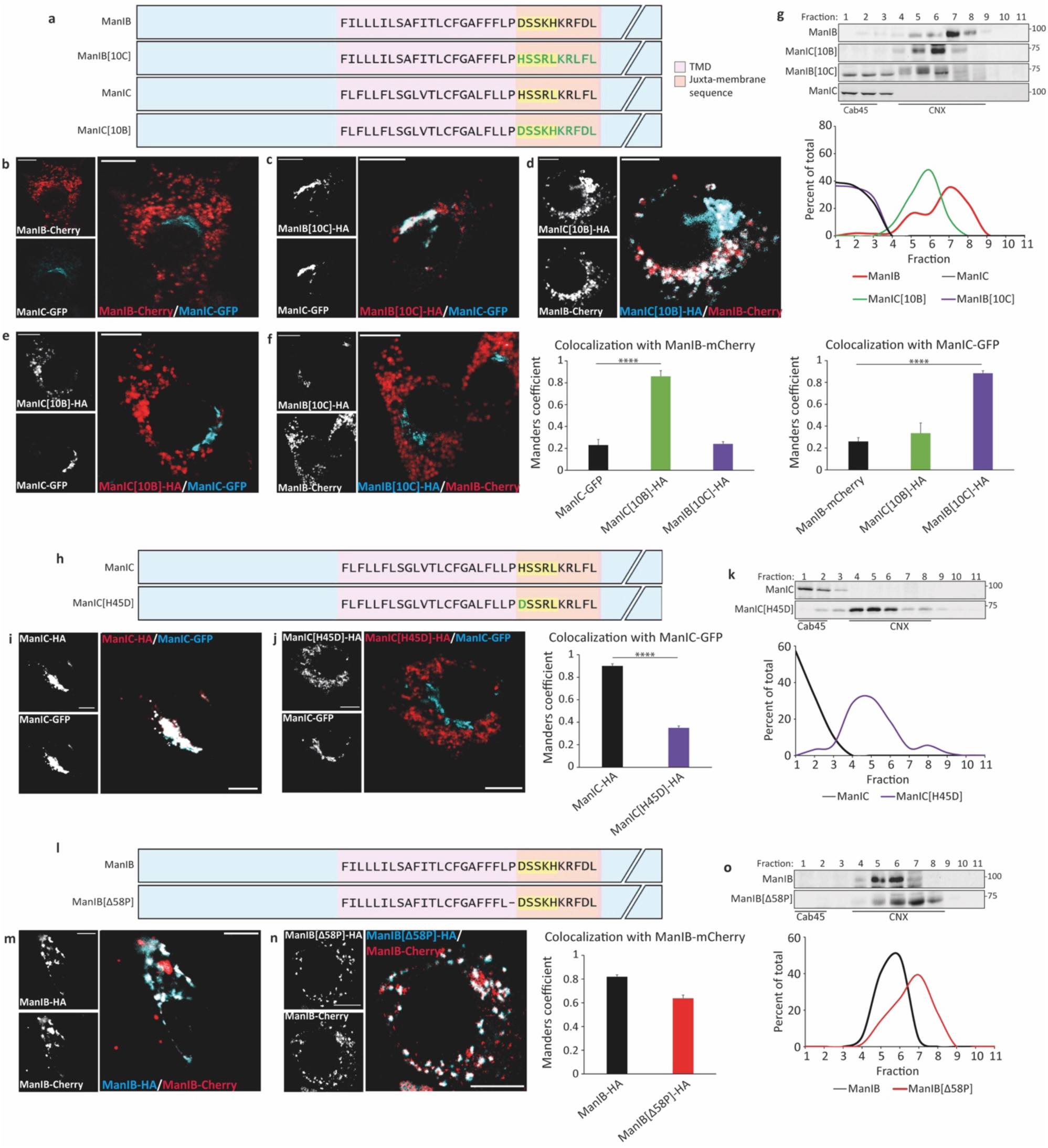
The juxtamembrane luminal peptide and its charges determine the QCV or Golgi localization of ManIB and ManIC. **(a, h, j)** Schemes illustrating the sequences in the transmembrane and juxta-membrane regions of ManIB and ManIC, and the mutants with the juxtamembrane luminal segment swapped ManIB[10C] and ManIC[10B] (a), or the mutants ManIC[H45D] (h) and ManIB[Δ58P] (j). **(b-f)** NIH 3T3 cells were transfected with (b) ManIB-mCherry and ManIC-GFP or (c) ManIB[10C]-HA and ManIC-GFP or (d) ManIC[10B]-HA and ManIB-mCherry or (e) ManIC[10B]-HA and ManIC-GFP or (f) ManIB[10C]-HA and ManIB-mCherry. 24 hours post transfection the cells were fixed and stained as in Fig. 5, using goat anti-mouse IgG conjugated to Dylight594 (c, e) or to Dylight488 (d, f) and were imaged on a confocal microscope. Bars=10μM. The graphs represent the colocalization of ManIC-GFP and each mutant with ManIB-mCherry (left, n=15, *P* value = 2.84 · 10^−11^) or ManIB-mCherry and each mutant with ManIC-GFP (right, n=15, *P* value = 5.34 · 10^−11^) as inferred by Mander’s coefficient. **(g, k, o)** HEK 293 cells were transfected with plasmids as indicated on the left of each panel (WT tagged with GFP and mutants tagged with HA). The cells were homogenized, and the homogenates run on iodixanol gradients as in Fig. 3 and plotted as a percent of the total. **(i, j, m, n)** Same as (b-f) with cells transfected with (i) ManIC-HA and ManIC-GFP or (j) ManIC[H45D]-HA and ManIC-GFP or (m) ManIB-HA and ManIB-mCherry or ManIB[Δ58P]-HA and ManIB-mCherry. The graphs represent the colocalization of ManIC-HA or ManIC[H45D]-HA with ManIC-GFP (n=13, *P* value = 4.76 · 10^−9^) or ManIB-HA and ManIB[Δ58P]-HA with ManIB-mCherry (n=15) as inferred by Mander’s coefficient.

It is noteworthy that a proline is present in the sequences of ManIA, ManIB and ManIC, at the end of the TMD, but not in ERManI (Fig. 2). As proline is a helix breaker, we wondered if its removal would have any consequences. We made a construct where proline at position 58 of ManIB was deleted. This change had almost no effect on the localization of the mutant ManIB[Δ58P] (Fig. 6l-o), suggesting that absolute continuity of the TMD helix in the luminal interface is not required for QCV localization.

Altogether, these results suggest that the distribution and orientation of charges at the transmembrane-lumen interface functions as a localization signal of these enzymes at either QCVs or at the Golgi complex.

### Alteration of charge distribution relocates GALT from the Golgi to QCVs

To test whether the juxtamembrane localization signal is sufficient to relocate an unrelated protein, we made mutants of β-1,3 galactosyltransferase (GalT), a type II membrane glycoprotein, anchored to the Golgi membrane, with a galactosyltransferase catalytic domain facing the lumen. We replaced the luminal juxtamembrane pentapeptide of GalT with that of ERManI or exchanged the positively charged arginine in position 1 after the TMD of GalT for an uncharged alanine residue, generating the mutants GalTER and GalT[R45A] respectively (Fig. 7a, Table 1). GalT contains an additional positive charge in one helical face compared to ERManI, which is removed in both mutants, achieving a similar charge pattern and orientation in this face between the two proteins (Fig. 7b). The fluorescent protein does not appear to influence the localization, as GalT-YFP colocalized almost completely with GalT-CFP (Fig. 7c). When observing the mutants in live-cell fluorescence microscopy, they appear in a punctate pattern, sharing a high level of colocalization with ERManI-mCherry and not colocalizing with GalT-CFP (Fig. 7d-i). In iodixanol equilibrium sedimentation gradients, GalT-YFP migrated in the lightest fractions, as expected, while GalTER and GalT[R45A] mutants shifted to different degrees to the heavier density ER/ QCV fractions (Fig. 7j). These results suggest that the luminal juxtamembrane charged pentapeptide of GalT is essential for its Golgi localization, which is changed to a punctate QCV pattern when altering the distribution of charges in this motif.

**Fig.7.**
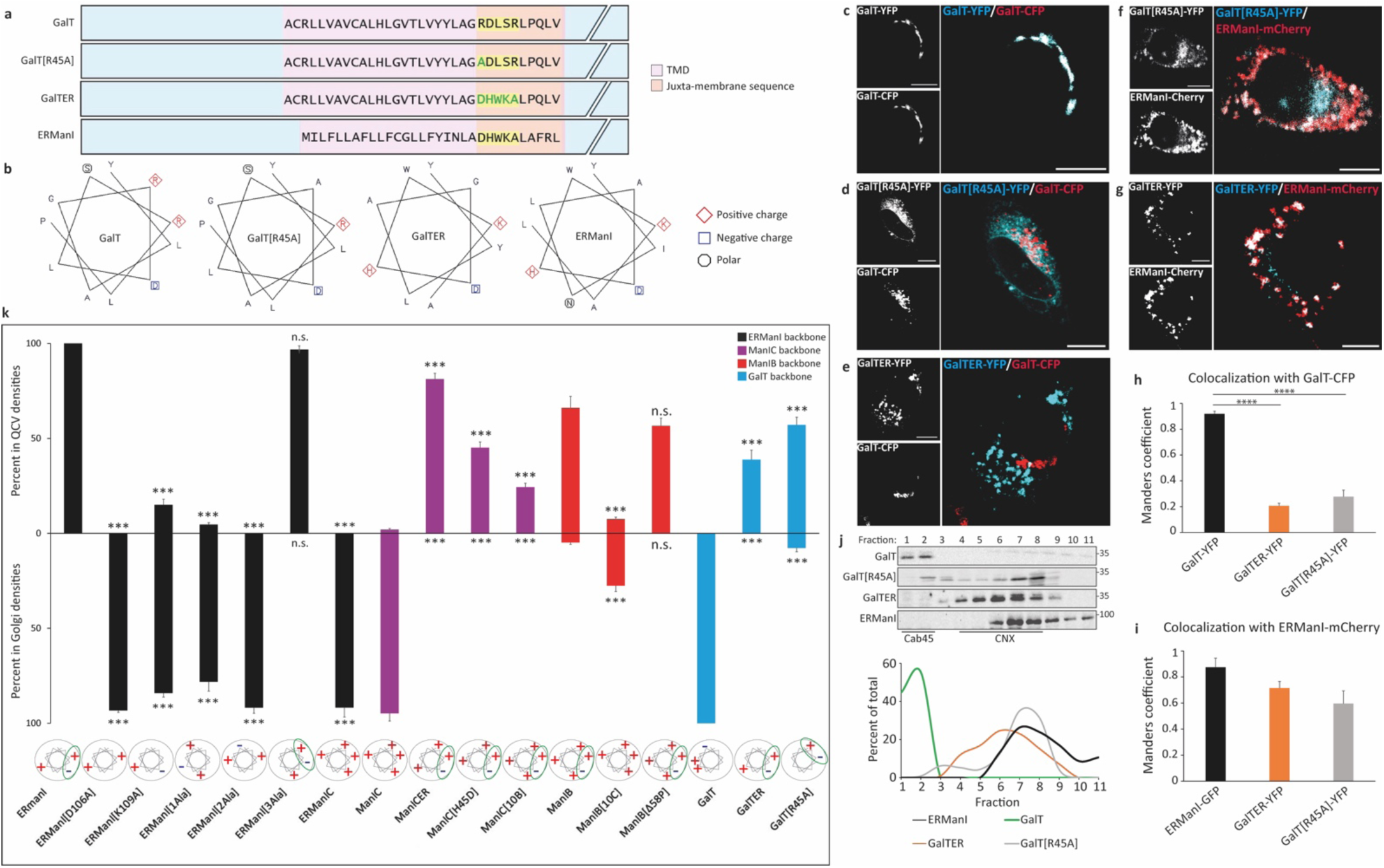
The juxtamembrane luminal peptide of ERManI or its charge distribution can be transferred to GalT, shifting its localization from the Golgi to QCVs. **(a)** Schemes illustrating the sequences in the transmembrane and juxtamembrane regions of GalT, ERManI and the mutants GalT[R45A] and GalTER, with the juxtamembrane luminal segment swapped with ERManI. **(b)** Edmonson alpha helix wheels of the luminal-transmembrane interfaces of GalT and its mutants compared to ERManI. **(c-g)** NIH 3T3 cells were transfected with plasmids expressing GalT-CFP together with **(c)** GalT-YFP or **(d)** GalT[R45A]-YFP or **(e)** GalTER-YFP or ERManI-mCherry together with **(f)** GalT[R45A]-YFP or with **(g)** GalTER-YFP and observed live at 24 hours post transfection. GalT-CFP has been pseudo-colored red. Bars=10μM. **(h, i)** Graph representation of colocalization of each transfected construct with **(h)** GalT-CFP (n=14, *P* value (GalTER-YFP) = 3.15 · 10^−11^, *P* value (GalT[R45A]-YFP) = 4.27 · 10^−11^), or **(i)** ERManI-mCherry (n=13, *P* value (GalT[R45A]-YFP) = 0.0394) as inferred by Mander’s coefficient. **(j)** HEK 293 cells were transfected with plasmids expressing GalT-YFP, GalT[R45A]-YFP, GalTER-YFP or ERManI-GFP. The cells were homogenized, and the homogenates run on iodixanol gradients as in Fig. 3 and plotted as a percent of the total. **(k)** QCV or Golgi localization of the different WT and mutant mannosidases as determined by the density gradients. The backbone of each protein is indicated in a different color. The green circles highlight the charge pattern in the Edmonson helix wheels at the bottom that signal QCV localization.

Altogether the results, summarized in Fig. 7k, clearly suggest the existence of a short, charged peptide at the TMD-luminal interface of these type II transmembrane proteins, with its charge distribution and 3D orientation determining QCV or Golgi subcellular localization.

## Discussion

Most studies on transmembrane protein localization have focused on signals within the cytosolic domains or TMDs ^39–41^. In contrast, our work demonstrates that the charge distribution and spatial orientation of residues at the TMD-luminal interface are critical determinants of subcellular localization for several type II transmembrane glycoproteins—ERManI, ManIB, ManIC, and GalT.

Using site-directed mutational analysis (Table 1), we show that altering either the distribution or orientation of charged residues at this interface significantly affects protein localization. For instance, substitution of charged residues with an uncharged alanine relocated ERManI, which normally resides in QCVs, to the Golgi apparatus, as can be observed in fluorescent microscopy of the mutants ERManI[D106A] and ERManI[K109A] (Fig. 3b-f and Suppl. Fig. S3). This relocalization was confirmed biochemically by iodixanol gradient sedimentation, where the migration of these mutants shifted to the lighter fractions, while WT ERManI migrated in the denser fractions (Fig. 3g, h). Conversely, the Golgi localization of ManIC was shifted to QCVs by a change of a positive histidine to a negative aspartate at the interface in ManIC[H45D] (Fig. 6h-k). On the other hand, there was a dramatic effect of insertion of alanine residues at the interface in ERManI (Fig. 4), without changing the net charge. Altogether, these results suggest that QCV localization is dictated not simply by net charge, but by a specific spatial pattern of charged residues at the TMD-luminal interface (Fig. 7k).

AlphaFold predicts with high confidence a helical structure in this region for ERManI but with less confidence for the other mannosidases. One possibility was that this helical extension might contribute to the localization of ERManI by extending the TMD. However, the alanine insertion mutants suggest that this is not the case. Although alanine promotes helix formation ^42, 43^, the insertion of one or two residues (ERManI[1Ala], ERManI[2Ala]) changed the localization to the Golgi, while insertion of three alanines (ERManI[3Ala]) restored the original QCV localization. These results suggest that helical pitch shifts—altering the orientation of the charged face relative to the TMD—are the key factor. Inserting three alanines shifts the helix by ∼300°, restoring the original orientation of the charged peptide on one face of the helix respect to the TMD (Fig. 4). Thus, localization appears to depend on the three-dimensional spatial orientation of key residues rather than their presence or charge alone.

The requirement for a specific orientation on one face of the helix raises the possibility that an adaptor protein may recognize both the TMD and the luminal juxtamembrane motif to promote the QCV localization. Such a protein would likely require a specific helical face presentation for binding, suggesting a mechanism involving protein-protein recognition. This is an interesting possibility that should be explored in future studies. The similarity of the charge pattern of ERManI with that of the determinant for ERQC localization of the ERAD substrate H2a is noteworthy (Fig. 1). This peptide, EGHRG, impedes the exit of H2a to the Golgi and from there to the cell surface ^15, 16^. This can be overcome by assembly of H2a with a mutated H1 subunit of the ASGPR, where the same charged peptide was inserted ^44^, implying that the putative adaptor protein could be displaced by specific interactions with another protein partner.

Mutational analysis of ManIB and ManIC further supports the role of this TMD-lumen interface as a conserved localization signal. Reciprocal exchanges of their juxtamembrane regions reversed their localization: ManIB[10C] shifted to the Golgi, while ManIC[10B] relocated to QCVs (Fig. 6b-g). Similar results were obtained with the mutants ManICER and ERManIC, which appeared in a similar pattern to that of ERManI (QCV) and ManIC (Golgi), respectively (Fig. 5). The dramatic change in localization of the mutant ManIC[H45D] reinforces the role of charge distribution as a determinant of localization (Fig. 6i-k). Interestingly, the removal of proline from the TMD-luminal interface of ManIB had almost no influence on the localization of the mutant ManIB[Δ58P] (Fig. 6m-o), implying that helix continuity (as in ERManI) is not required to achieve the proper protein localization. Contrary to alanine, the presence of proline causes only a minor shift of approximately 40° in the helical pitch ^45, 46^, which does not significantly change the orientation of the charged peptide with respect to the TMD.

To test the generality of this mechanism, we examined the Golgi-resident galactosyltransferase. Substitution of the juxtamembrane peptide of GalT with that of ERManI led to a redirection of the mutant GalTER from the Golgi to QCVs (Fig. 7e, g-j). Neutralization of the additional positive charge in the juxtamembrane luminal peptide of GalT resulted in a similar change in localization of the mutant GalT[R45A], strengthening the notion that the TMD-luminal interface determinant may function as a universal signal for organelle-specific localization in type II transmembrane proteins. (Fig. 7). Similar universality has been previously described by Bonifacino and Glick, who proposed that vesicular trafficking depends on highly conserved cytosolic sorting signals ^47^.

Together, our data establish the TMD-lumen interface as a novel localization signal based on three-dimensional arrangement of charges. This opens new directions for studying protein-lipid and protein-protein interactions through this interface, that govern intracellular sorting. Further work should aim to identify the putative adaptor proteins or cofactors that interpret these spatial signals.

In a broader context, mislocalization of membrane proteins plays a role in numerous diseases ^48, 49^. While mutations affecting charge distribution have been studied for their impact on protein topology, structure or interactions ^50^, their direct effect on intracellular targeting has received less attention. Our identification of a localization signal based on the 3D arrangement of charged residues at the TMD-lumen interface offers a new framework for understanding how specific topological features determine protein trafficking and organelle targeting, potentially leading to therapeutic strategies that restore proper localization by manipulating these motifs.

## Acknowledgements

We would like to thank A. Herscovics and K. Hirshberg for plasmids and M. Schushan and N. Ben Tal for help with the bioinformatics survey. Work was supported by grant 2577/20 from the Israel Science Foundation (GZL).

## Author Contributions

Conceptualization: GZL. Investigation: HS, RB, TM. Supervision: GZL. Writing: HS, GZL. Review and editing: HS, RB, TM, GZL.

