## Supplementary material for "Helical charge distribution at the transmembrane-luminal interface determines subcellular localization": Suppl.

### Supplementary information

### Supplementary figures

**a** ERManI

| | |
|---|---|
| Homo sapiens (human): | KQLSRLQRNMILFLLAFLLFCGLLFYINLADHWKALAFRLEEEQK |
| Pan troglodytes (chimpanzee): | KQLSRLQRNMILFLLAFLLFCGLLFYINLADHWKALAFRLEEQQK |
| Macaca mulatta (rhesus): | KQLSRLQRNMILFLLAFLLFCGLLFYINLADHWKALAFRLEEEQK |
| Canis familiaris (dog): | KQLSRLQRNVILFSLTFLMLCGFLSYISVADQWTAVDGRSAEEQK |
| Rattus norvegicus (Norwegian rat): | KQLSRLQRNVILFVLGFLILCGFLYSLQVSDQWKALSGSRAEVEQ |
| Mus musculus (mouse): | KQLSRLQRNVILFVLGFLILCGFLYSLHTADQWKALSGRPAEVEK |
| Equus caballus (horse): | KQLSRLQRNVILFLLAFLLLCGLLSYISVADQWTAPAGRSAEEHK |
| Gallus gallus (Chicken): | KQLSRLQRSIILFLFAFLTVCGVISYTSVREPWKSLTSKSSDEHG |
| Monodelphis domestica (opossum): | KQLSRLQRNIILFFFAFLAVCGLISYINMANHGRAFTSRSSEDQK |
| Xenopus tropicalis (pipid frog): | KQLSRLQRNIILFLCVLLAVCAVGSYSNLGEHWRALTSQSEEHYE |
| Xenopus laevis (African clawed frog): | KQLSRFQRNIILFLCVVLAVCTVVSFSNLGEHWRALISQSEEHYE |
| Danio rerio (zebrafish): | KQLSRLQRSLILFLLAFLFICGILSYSSLTEQWRGISDRSLKEDW |
| Tetraodon nigroviridis (Green spotted pufferfish): | KQLSRLQRSLILFLLMLLLIFGLFSFPTITEQWKG |
| Oryzias latipes (fish): | KQLSQLQRSLILFILALLFICGIASFPTFNEHLRGLVFGRDRREF |
| Takifugu rubripes (fish): | KQLSRLQRSLILFLLMLLLIFGLFSFPSITEQWRGGKSLDQAAEKD |
| Gasteroteus aculeatus (fish): | KQLSKLQRSLILLVLVICGMASYPIVTEHLRGLFLQQKSV |

**b** ManIA

| | |
|---|---|
| Homo sapiens (human): | SGPAALRLTEKFVLLLVFSAFITLCFGAIFFLPDSSKLLSGVLFHSSPAL |
| Mus musculus (mouse): | SGPAALRLTEKFVLLVFSAFITLCFGAIFFLPDSSKLLSGVLFHSNPAL |
| Rattus norvegicus (Norwegian rat): | SGPAALRLTEKVLLLVFSAFITLCFGAIFFLPDSSKLLSGVLFHSNPAL |
| Bos taurus (cattle): | SGPAALRLTEKFVLLLVFSAFILTLCFGAIFFLPDSSKLLSGVLFHSSPAL |

**c** ManIB

| | |
|---|---|
| Homo sapiens (human): | PPSFPHHRATLRLSEKFILLLILSAFITLCFGAFFFLPDSSKHKRFDLGLEDVLI |
| Pan troglodytes (chimpanzee): | PPSFPHHRATLRLSEKFILLLILSAFITLCFGAFFFLPDSSKHKRFDLGLEDVLI |
| Rattus norvegicus (Norwegian rat): | PPSFPHHRATLRLSEKFILLLILSAFITLCFGAFFFLPDSSKHKRFDLGLEDVLI |
| Mus musculus (mouse): | PPSFPHHRATLRLSEKFILLLILSAFITLCFGAFFFLPDSSKHKRFDLGLEDVLI |
| Xenopus laevis (African clawed frog): | TSSFPHHRATLRLSEKFILLLILSAFITLCFGAFFFLPDSSKHKRFDLGLEDVLI |
| Sus scrofa (pig): | PPSFPHHRATEKFILLLILSAFITLCFGAFFFLPDSSKHKRFDLGLEDVLI |
| Tetraodon nigroviridis (Green spotted pufferfish): | PHHRATLRLSEKFILLLILSAFITLCFGAFFFLPDSSKHKRFDLGLEDVLI |
| Spodoptera frugiperda (fall armyworm): | VPSISRRSFRLREKYLIVSVLLTFGIVWLGALFYLPEFKSSNSVNDSVYNVYK |

**d** ManIC

| | |
|---|---|
| Homo sapiens (human): | PGFVPASPWGLRLPQKFLFLLFLSGLVTLCFGALFLLPHSSRLKRLFLAPRTQQP |
| Equus caballus (horse): | PGFVPASPWGLRLPQKFLFLLFLSGLITLCFGALFLLPKSSRLKRLFLAPRTQQP |
| Rattus norvegicus (Norwegian rat): | PGFVPASSWGLRLPQKFLFLLFLSGLVTLCFGALFLLPHSSRLKRLFSAPRTQQP |
| Mus musculus (mouse): | PGFVPASPWGLRLPQKFLFLLFLSGLVTLCFGALFLLPHSSRLKRLFSAPRTQQP |

**Supplementary Fig. S1. Sequence comparison of orthologs of ERManI and mannosidases IA, IB and IC.** Predicted transmembrane region is underlined. Positively charged amino acids are in red and negatively charged in blue. The juxtamembrane luminal pentapeptide with conserved charge, except in ManIC, is highlighted in yellow.

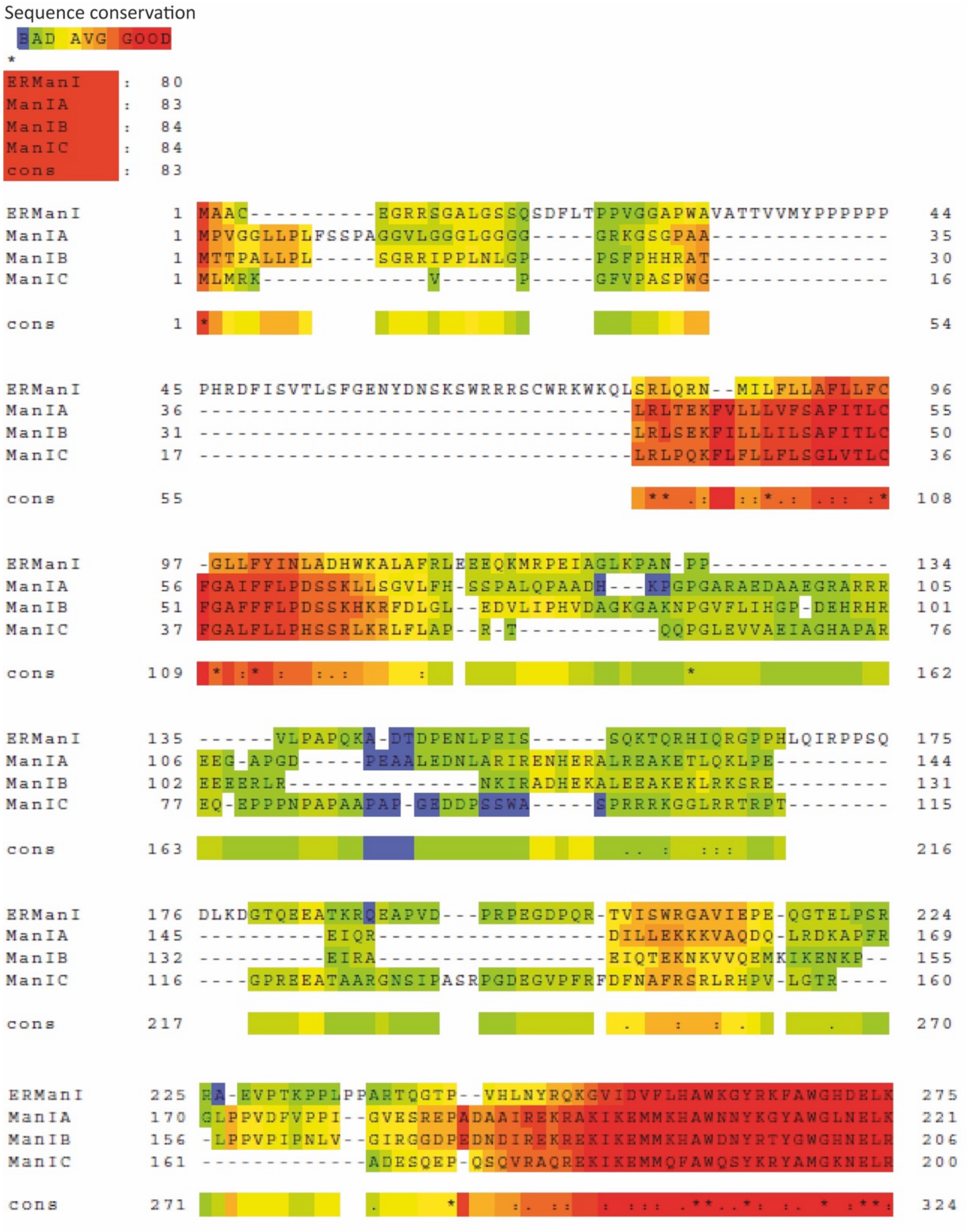

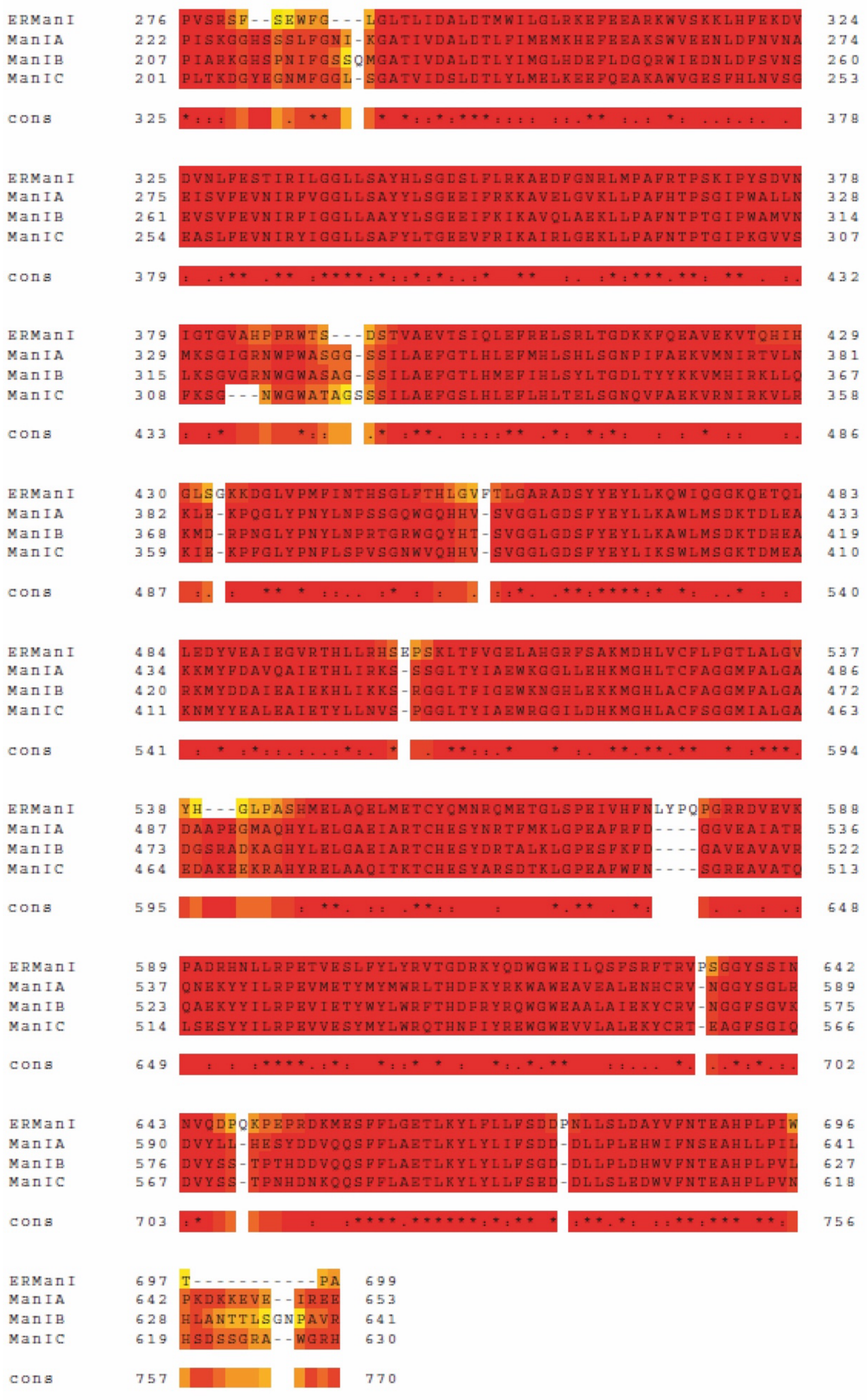

**Supplementary Fig. S2. Sequence alignment of ERManI, ManIA, ManIB and ManIC.** Sequence alignment of ERManI, ManIA, ManIB and ManIC performed by T-COFFEE software based on combination of several algorithms including Clustal W. The two most conserved regions highlighted in red correspond to the TMD near the N terminus and the long catalytic domain near the C terminus.

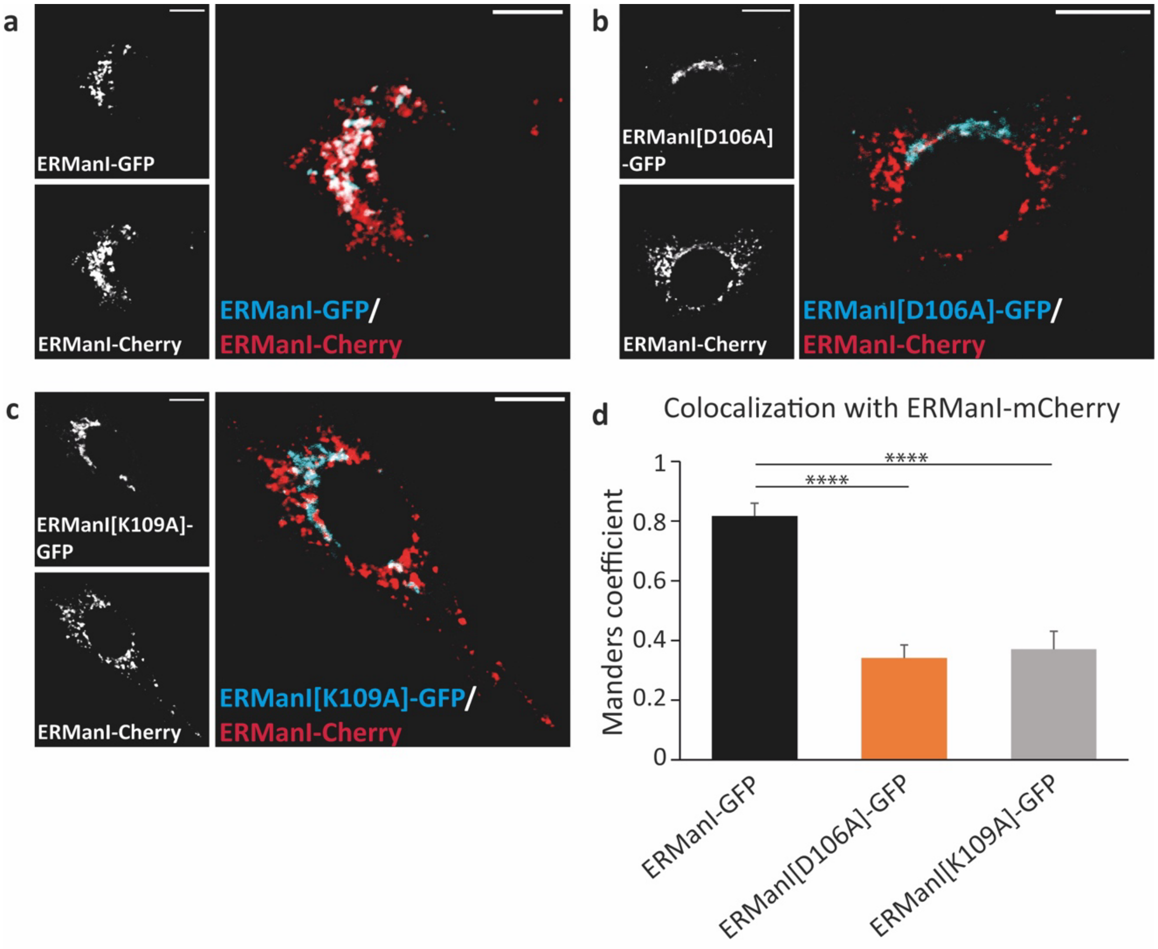


**Supplementary Fig. S3. Fluorescence of ERManI and charge mutants in fixed but unpermeabilized cells. (a-c)** NIH 3T3 cells were fixed by 3% PFA and left unpermeabilized, 24 hours following transfection with ERManI-mCherry and either **(a)** ERManI-GFP or **(b)** ERManI[D106A]-GFP or **(c)** ERMan1[K109A]-GFP. Bars=10mM. **(d)** Graph representation of colocalization of each transfected construct with ERManI-cherry as inferred by Manders coefficient (n=10, *P* value (ERManI[D106A]-GFP) = $2.16 \cdot 10^{-5}$, *P* value (ERManI[K109A]-GFP) = $2.41 \cdot 10^{-8}$).

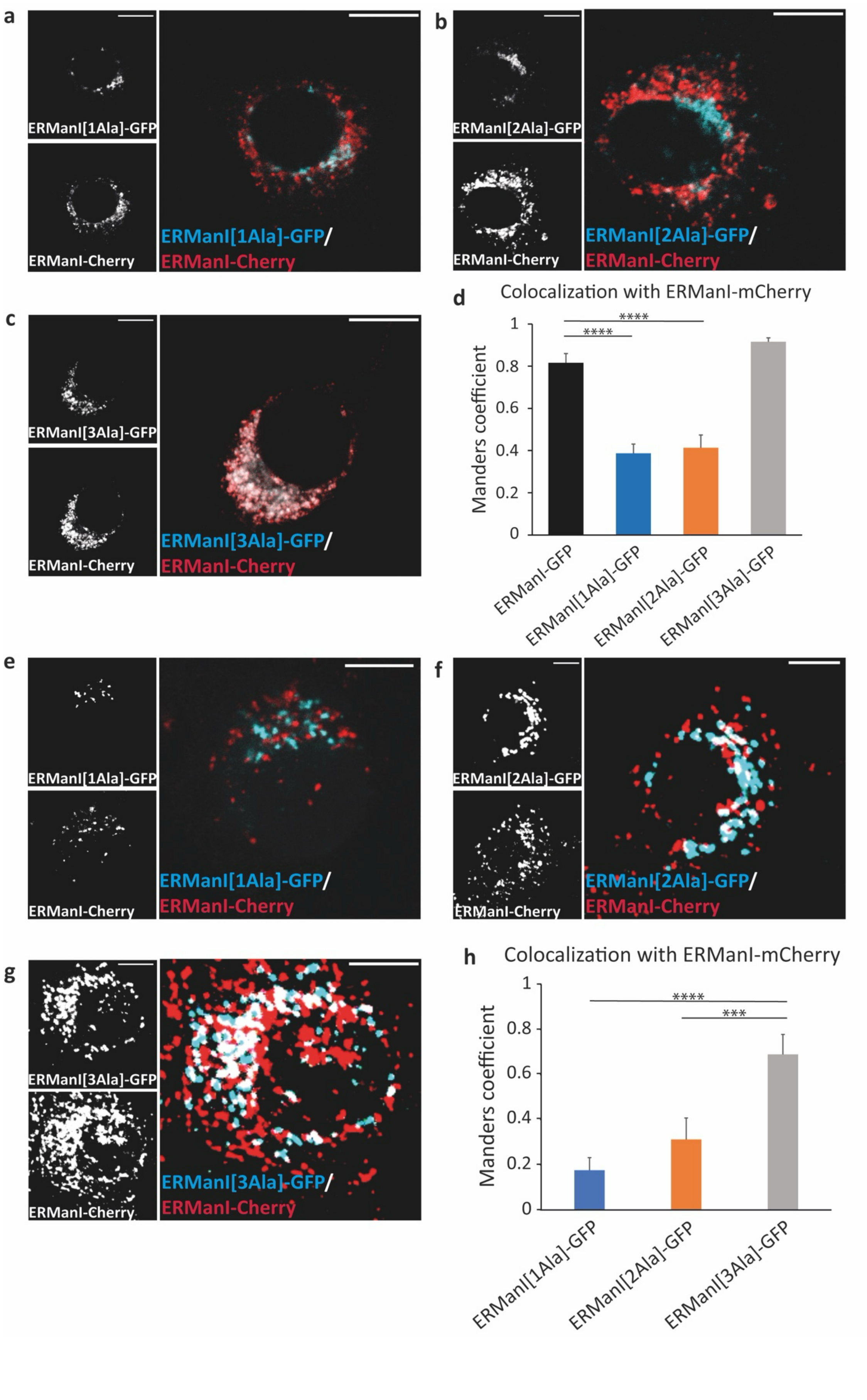
a
ERManI[1Ala]-GFP
ERManI-Cherry
ERManI[1Ala]-GFP/
ERManI-Cherry
b
ERManI[2Ala]-GFP
ERManI-Cherry
ERManI[2Ala]-GFP/
ERManI-Cherry
c
ERManI[3Ala]-GFP
ERManI-Cherry
ERManI[3Ala]-GFP/
ERManI-Cherry
d
Colocalization with ERManI-mCherry
Manders coefficient
0
0.2
0.4
0.6
0.8
1
****
****
ERManI-GFP
ERManI[1Ala]-GFP
ERManI[2Ala]-GFP
ERManI[3Ala]-GFP
e
ERManI[1Ala]-GFP
ERManI-Cherry
ERManI[1Ala]-GFP/
ERManI-Cherry
f
ERManI[2Ala]-GFP
ERManI-Cherry
ERManI[2Ala]-GFP/
ERManI-Cherry
g
ERManI[3Ala]-GFP
ERManI-Cherry
ERManI[3Ala]-GFP/
ERManI-Cherry
h
Colocalization with ERManI-mCherry
Manders coefficient
0
0.2
0.4
0.6
0.8
1
****
***
ERManI[1Ala]-GFP
ERManI[2Ala]-GFP
ERManI[3Ala]-GFP

**Supplementary Fig. S4. . Fluorescence of ERManI and alanine insertion mutants. (a-d)** NIH 3T3 cells were fixed by 3% PFA and left unpermeabilized, 24 hours following transfection with ERManI-mCherry and either **(a)** ERManI[1Ala]-GFP or **(b)** ERManI[2Ala]-GFP or **(c)** ERManI[1Ala]-GFP. Bars=10mM. **(d)** Graph representation of colocalization of each transfected construct with ERManI-cherry as inferred by Manders coefficient. (n=10, *P* value (ERMan1ala-GFP) = $8.32 \cdot 10^{-6}$ , *P* value (ERMan2ala-GFP) = $4.56 \cdot 10^{-5}$. **(e-f)** U2OS cells were transfected with plasmids expressing ERManI-mCherry and **(e)** ERManI[1Ala]-GFP or **(f)** ERManI[2Ala]-GFP or **(g)** ERManI[3Ala]-GFP and observed live at 24 hours post transfection. Bars=10mM. **(h)** Graph representation of colocalization of each transfected construct with ERManI-cherry as inferred by Manders coefficient. (n=7, *P* value (ERMan1ala-GFP) = $4.211 \cdot 10^{-6}$ , *P* value (ERMan2ala-GFP) = $1.9 \cdot 10^{-4}$.
